# MPS1 localizes to microtubule-attached kinetochores and actively promotes microtubule release

**DOI:** 10.1101/2022.05.23.493048

**Authors:** Daniel Hayward, Emile Roberts, Ulrike Gruneberg

## Abstract

**Summary:** In eukaryotes, the spindle assembly checkpoint protects genome stability in mitosis by preventing anaphase onset until incorrect microtubule-kinetochore attachment geometries have been eliminated and chromosome biorientation has been completed. These error correction and checkpoint processes are linked by two conserved serine/threonine kinases, Aurora B and MPS1 ^1, 2^. In the prevailing model for spindle checkpoint signaling, MPS1 detects microtubule-free kinetochores generated by the Aurora B-dependent error correction pathway, and initiates spindle checkpoint signaling. However, we find that MPS1 initially localizes to microtubule-attached kinetochores in a manner dependent on the relative activities of Aurora B and its counteracting phosphatase PP2A-B56, and then actively promotes microtubule release. Thus, MPS1 is not a passive sensor for the microtubule binding state of kinetochores but actively generates microtubule-free checkpoint signaling kinetochores. MPS1 is thus instrumental for both the initial resolution of incorrect microtubule-kinetochore attachments and the downstream propagation of spindle checkpoint signaling.

## Main text

Aurora B initiates the removal of erroneous microtubule-kinetochore attachments and hence generates unattached kinetochores ^1, 3-7^. In the canonical model for error correction, MPS1 is then recruited to the microtubule-free kinetochores generated by Aurora B and catalyzes the assembly of the diffusible mitotic checkpoint complex, MCC ^8-10^. MCC globally inhibits the anaphase promoting complex/cyclosome ubiquitin E3 ligase necessary for anaphase onset ^11^. MPS1 activity thus links the presence of unattached kinetochores to the arrest of mitotic progression. MPS1 and microtubules are thought to compete for binding sites at the outer kinetochore, with incoming microtubules replacing MPS1 during spindle assembly to trigger checkpoint silencing ^12, 13^. However, not all published work is consistent with this model. Checkpoint silencing has been shown to occur well below full occupancy of microtubule binding-sites at the kinetochore, a competition model therefore cannot completely explain the observed loss of MPS1 from the entire kinetochore upon microtubule attachment and the switch-like change of kinetochore status from checkpoint-active to checkpoint-inactive ^14, 15^. Furthermore, MPS1 is required for chromosome biorientation and correction of microtubule-kinetochore attachments errors ^16-20^. This is not readily explained within the constraints of the current model where MPS1 localizes exclusively to unattached kinetochores generated by the error correction process. Therefore, it is not clear how MPS1 would exert a function in the correction of erroneous microtubule-attachment geometries, which must require direct interaction with microtubule-attached kinetochores.

We therefore explored whether MPS1 recruitment to kinetochores is coupled to microtubule occupancy through kinase and phosphatase activities regulating the “*state*” of the kinetochore, rather than through direct competition for binding sites. Two lines of evidence support this idea. First, the centromeric Aurora B kinase is known to promote MPS1 kinetochore localization directly, independently from its role in generating microtubule-free “unattached” kinetochores ^3-7, 21^. Second, the balance between the kinase activity of Aurora B and the opposing phosphatase PP2A-B56, respectively, is critical for MPS1 binding and release at unattached kinetochores ^21^. Such a model predicts a transient transition state in which MPS1 and microtubules are bound to kinetochores simultaneously. First, to investigate this idea, it was important to ascertain that under normal growth conditions microtubule-attached kinetochores exhibiting high levels of MPS1 could be observed. As predicted, in prometaphase cells, kinetochores staining positive for both the attached-kinetochore marker astrin ^22^, as well as for MPS1, were present in small numbers (Extended Data Fig. 1A-D). Stability at reduced temperature has been widely used to distinguish stable and transient microtubule-kinetochore attachments, respectively ^23^. We therefore tested if, by altering the balance of kinase and phosphatase activity, reduced temperature could stabilize the normally transient state in which MPS1 and microtubules are bound simultaneously to the kinetochore (Fig. 1A). Brief cold treatment significantly increased the number of attached kinetochores exhibiting high levels of MPS1, equivalent to unattached kinetochores, at metaphase plates (Fig. 1B and C). To facilitate the direct comparison of microtubule-attached and unattached states in the same cell, we performed the same analysis in HeLa cells treated with the KIF11/Eg5 inhibitor S-Trityl-L-cysteine (STLC) to create monopolar spindles. Kinetochore pairs in monopolar spindles typically contain one attached and one unattached kinetochore, and cold treatment resulted in an elevated frequency of kinetochores displaying equivalent MPS1 levels on both kinetochores (Fig. 1D and E, Extended Data Fig. 1E-G). Importantly, microtubule-kinetochore attachments were not changed under these conditions (Extended Data Fig. 1H-K).

**Fig. 1.**
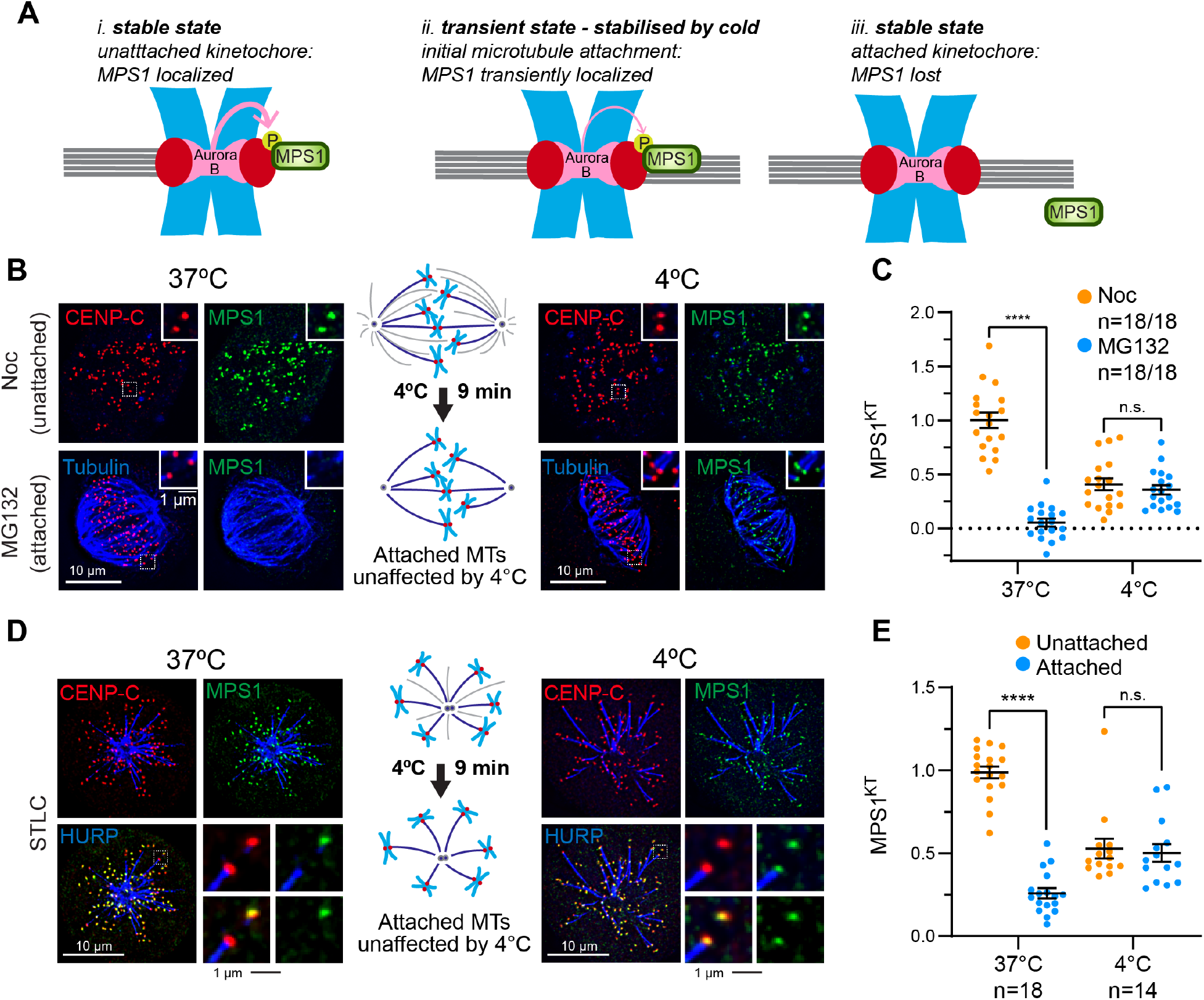
MPS1 can localize to microtubule-attached kinetochores. (**A**) Schematic diagram depicting transient MPS1 binding to microtubule-attached kinetochores. (**B**) Nocodazole (Noc) or MG132 arrested HeLa MPS1-GFP cells were either directly fixed or cooled to 4°C for 9 min and then fixed, and immuno-stained as indicated. (**C**) MPS1 kinetochore cell average intensities of cells in (B). Bars show mean ± S.E.M. (**D**) STLC-arrested HeLa MPS1-GFP-cells were treated as in (B). The K-fibre marker HURP was used to visualize kinetochore attachments ^45^. (**E**) MPS1 kinetochore cell average intensities of cells in (D).

Prompted by our observation that the phosphatase PP2A-B56 regulates MPS1 levels at unattached kinetochores ^21^, we then asked if inhibition of specific phosphatase activities would stabilize MPS1 localization to attached kinetochores. To test this idea, we examined endogenous MPS1 localization in HeLa MPS1-GFP cells ^24^ briefly treated with the potent PP2A and PP1 inhibitor calyculin A. Compared to untreated cells, this treatment significantly increased the amount of MPS1 detected at bioriented kinetochores in the metaphase plate without altering microtubule-kinetochore attachments (Fig. 2A and B, Extended Data Fig. 2A and B). Previous work has established that PP2A-B56 is recruited to kinetochores via association with the BUBR1 spindle checkpoint protein ^25, 26^. To test the involvement of this specific pool of PP2A-B56, we replaced endogenous BUBR1 with the PP2A-B56 binding deficient BUBR1-L669A/I672A mutant (BUBR1^LI/AA^) using an RNAi rescue strategy (Extended Data Fig. 3A) ^25, 27^. BUBR1 replacement with the B56-binding deficient GFP-BUBR1^LI/AA^ in STLC-treated monopolar spindles reduced the number of microtubule-attached kinetochores (Extended Data Fig. 3B-D) but in kinetochore pairs with a clear attachment, MPS1 was recruited to both the attached and the unattached kinetochore, as also observed for calyculin A treatment (Fig. 2C-D, Extended Data Fig. 2C-F). This provides evidence for the existence of a normally transient kinetochore state where microtubules and MPS1 are simultaneously bound and which is limited by the subsequent MPS1-dependent recruitment of BUBR1-PP2A-B56 complexes. MPS1 therefore transiently localizes to microtubule-attached kinetochores, consistent with a model in which dynamic phosphorylation controls MPS1 localization in response to microtubule occupancy, and we set out to understand how this is regulated in more detail.

**Fig. 2.**
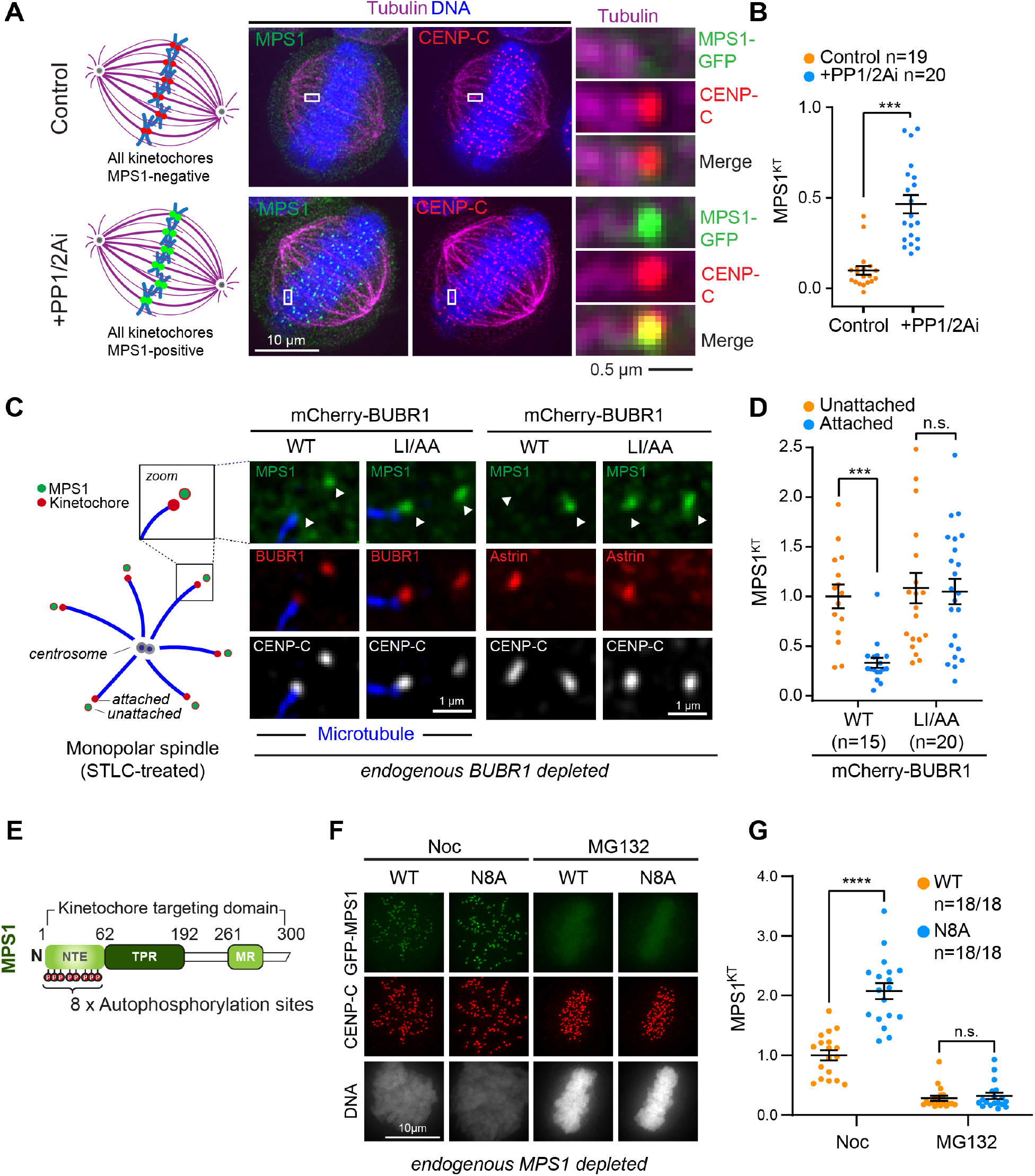
PP2A-B56 phosphatase activity is required to release MPS1 from attached kinetochores. (**A**) HeLa cells expressing MPS1-GFP were arrested with 20 μM MG132 for 2 hrs, treated with 25 nM calyculin A (PP1/2Ai) for 6 mins and immuno-stained as indicated. Representative attached kinetochores have been enlarged on the right. (**B**) Cell averages of kinetochore-MPS1 intensities at attached metaphase kinetochores in cells treated as in (A). Bars represent mean± S.E.M. Values were normalized to the levels of MPS1 at unattached kinetochores without PP1/2Ai treatment (see Extended Data Fig. 2A-B). (**C**) STLC arrested Flp-In TRex HeLa cells expressing GFP-MPS1 from the endogenous promoter were depleted of endogenous BUBR1 and induced to express mCherry-BUBR1^WT^ or PP2A-B56 binding deficient mCherry-BUBR1^LI/AA^. Cells were immuno-stained as indicated. Representative monotelic kinetochore pairs are shown (white arrowheads). (**D**) Cell averages of kinetochore-MPS1 at unattached or attached kinetochores in cells treated as in (C). Values were normalized to the mean of the unattached mCherry-BUBR1^WT^ control. Bars show mean± S.E.M. (**E**) Schematic diagram of the MPS1 N-terminal kinetochore targeting domain. Autophosphorylation sites are indicated as magenta lollipops. (**F**) Flp-In TRex HeLa cells depleted of endogenous MPS1 and expressing GFP-MPS1^WT^ or GFP-MPS1^N8A^ were treated with nocodazole/MG132 for 2 hrs (Noc) or just with MG132 (30 mins), and then fixed and stained as indicated. (**G**) Cell averages of MPS1 kinetochore intensities, normalized to the mean WT Noc value, were plotted. Bars indicate mean± S.E.M.

Auto-phosphorylation of the N-terminal kinetochore-binding domain of MPS1 has been suggested to modulate its localization at unattached kinetochores ^17, 28-30^. To test the role of auto-phosphorylation in localizing MPS1 to attached kinetochores, we generated GFP-tagged MPS1 mutants lacking all known autophosphorylation sites at S7, T12, T33 and S37 ^28^ and S15, T45, T46 and S49 ^31^ (Figure 2E), and used this version of MPS1, GFP-MPS1^N8A^, to replace the endogenous protein (Extended Data Fig. 4A). GFP-MPS1^N8A^ accumulated to 2-2.5 times higher levels than GFP-MPS1^WT^ at unattached kinetochores, (Fig. 2F and G, Extended Data Fig. 4B and C). This behavior was similar to kinase-dead MPS1 (GFP-MPS1^KD^) ^17, 30^. GFP-MPS1^N8A^ recruitment to kinetochores was not increased further by the addition of the MPS1 inhibitor AZ3146 ^17^, suggesting that all relevant autophosphorylation sites had been mutated (Extended Data Figure 4B and C). Corroborating these findings, phospho-mimetic mutations of the same sites (GFP-MPS1^N8E^) resulted in poor kinetochore localization which was not improved upon MPS1 inhibition (Extended Data Fig. 4B-C). If auto-phosphorylation of the MPS1 N-terminus was the key reason for loss of MPS1 from attached kinetochores, GFP-MPS1^N8A^ would be expected to remain localized to attached kinetochores in metaphase cells. However, GFP-MPS1^N8A^ was not observed on attached kinetochores at metaphase plates or in a monopolar spindle situation (Fig. 2F and G, Extended Data Fig. 4D and E). Therefore, we conclude that, although MPS1 N-terminal autophosphorylation does modulate the levels of MPS1 at unattached kinetochores, it is not the mechanism conferring sensitivity to microtubule attachment.

We next explored the role of Aurora B activity in MPS1 recruitment to microtubule-attached kinetochores. Inhibition of Aurora B activity with the small molecule inhibitor ZM447439 ^32^ results in the loss of MPS1 from unattached kinetochores, including the MPS1^KD^ (kinase-dead) and MPS1^N8A^ mutants which cannot be auto-phosphorylated ^17, 21^ (Extended Data Fig. 5A and B). To test whether Aurora B activity at kinetochores is not only necessary but also sufficient for MPS1 recruitment, we created a situation where Aurora B activity can be modulated at microtubule-attached kinetochores. To achieve this, we generated HeLa cells expressing both endogenously GFP-tagged MPS1 and a rapamycin-sensitive dimerization module which recruits a truncated version of the Aurora B interaction partner INCENP, together with Aurora B, to the outer kinetochore protein Mis12 upon rapamycin addition ^7, 33^ (Fig. 3A). These cells were then arrested at metaphase by proteasome inhibition with MG132, so all kinetochores were fully attached to microtubules, and MPS1 had left the kinetochore (Figure 3B; -1 min). Live cell imaging was used to analyze the behavior of MPS1-GFP upon rapamycin addition. When INCENP-Aurora B was targeted to attached kinetochores in this way, MPS1-GFP was rapidly recruited to microtubule-attached kinetochores, at levels which were at least equivalent to the levels of MPS1 observed at unattached prometaphase kinetochores (Fig. 3B and C and Extended Data Figure 6A-D). As a consequence of MPS1 recruitment to attached kinetochores, downstream spindle checkpoint proteins such as MAD1 were also detected (Extended Data Fig. 6C and E). In this situation, MPS1 recruitment to attached kinetochores was completely dependent on Aurora B activity (Figure 3D and E). Interestingly, microtubule attachment was maintained on average for 5 mins after rapamycin addition, after which time attachments started to be lost, and chromosomes left the metaphase plate (Fig. 3B and C). Cold treatment of cells 2 mins after rapamycin addition, confirmed that microtubule-kinetochore attachments were still intact at this point (Extended Data Fig. 6F and G) but were lost after longer incubation with rapamycin (Extended Data Figure 7). Therefore, MPS1 can be recruited to microtubule-attached kinetochores as long as Aurora B activity exceeds a threshold level. At microtubule-attached kinetochores, Aurora B activity is attenuated, and PP2A-B56 activity triggers loss of MPS1 from these kinetochores ^7, 21^ (Fig.2C and D). HEC1/NDC80 is the key microtubule binding component of the outer kinetochore and is thought to constitute the crucial Aurora B regulated binding site for MPS1 at checkpoint active, unattached kinetochores ^12, 34^. However, mutations of all Aurora phosphorylation sites in HEC1/NDC80 to alanine or phospho-mimetic aspartate did not reduce MPS1 recruitment or render it resistant to Aurora B inhibitors ^4^ (Extended Data Fig. 8). NDC80 is thus unlikely to be the crucial target for Aurora B in promoting recruitment of MPS1.

**Fig. 3.**
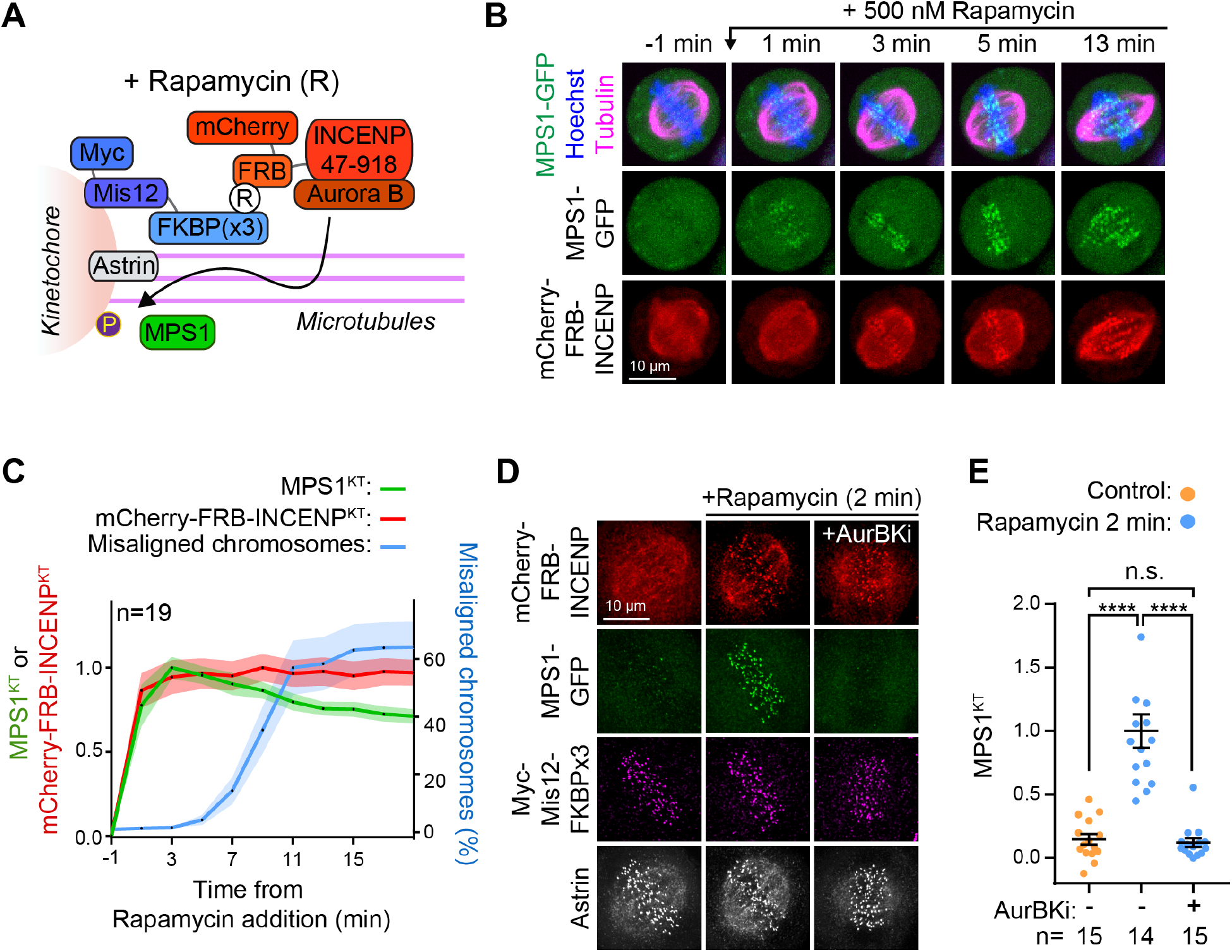
Aurora B-dependent localization of MPS1 to kinetochores can be uncoupled from microtubule-attachment status. (**A**) Diagram of the rapamycin-sensitive dimerization module used to recruit INCENP-Aurora B to the outer kinetochore protein Mis12. (**B**) Movie stills of a HeLa cell expressing Myc-MIS12-FKBPx3, mCherry-FRB-INCENP^47-918^ and MPS1-GFP, and incubated with SiR-tubulin and Hoechst. Cells were treated with MG132 30 min prior to imaging to enrich for cells with fully attached metaphase plates. Cell at -1 min is at metaphase prior to addition of 500 nM rapamycin. (**C**) Quantification of kinetochore intensities of MPS1-GFP (green), mCherry-FRB-INCENP (red) or misaligned chromosomes (proportion of Hoechst fluorescence signal outside of the metaphase plate) (blue) in cells as shown in (B). Line shows the mean, the shaded areas are S.E.M. (**D**) HeLa cells as in (B), arrested for 2 hours with MG132 (20 μM) and stained with anti-Myc and anti-Astrin. Aurora B inhibitor ZM447439 was added at 10 μM for 10 min prior to fixing, rapamycin was added at 500 nM 2 min prior to fixation. (**E**) Average MPS1 cell intensities, normalized against Myc-Mis12 intensities, at kinetochores in cells treated as in (D) were plotted. Bars represent mean± S.E.M.

The data presented so far suggest that MPS1-positive microtubule-attached kinetochore states are transient, either due to release of MPS1 or release of microtubules. MPS1 has been described as an important factor for error correction ^16-20^, yet how this activity is coordinated with Aurora B-mediated error correction has remained unclear. Our findings suggest that transient MPS1 localization to microtubule-attached kinetochores may be functionally important. To investigate the contribution of MPS1 to the resolution of microtubule-kinetochore attachments, we took advantage of our system for rapamycin-induced Aurora B recruitment to kinetochores, and repeated our experiments in the presence of MPS1 inhibitor. The kinetics of Aurora B-dependent MPS1 recruitment to attached kinetochores were not altered in this situation (Fig. 4A). However, MPS1 inhibition resulted in a marked delay in the loss of kinetochore attachments in comparison to control cells (Fig. 4A and B; Extended Data Fig. 7). This is consistent with a crucial role for MPS1 in the resolution of incorrect attachments. Our findings are in line with recent reports identifying Ska3 and HEC1/NDC80 as targets of MPS1 during error correction ^20, 35^. To confirm this role of MPS1 in the turn-over of erroneous microtubule-kinetochore attachments, high resolution imaging of MPS1 at syntelic attachments undergoing error correction in STLC-treated cells was carried out. This showed that MPS1 localization to microtubule-attached kinetochores precedes loss of attachment (Fig. 4C and D), confirming the idea that the presence of MPS1 at kinetochores is required for the prompt removal of the incorrect attachment. The current view of how spindle assembly checkpoint signaling is initiated therefore needs to be revised since these data show that MPS1 does not merely detect but actively generates unattached kinetochores. In this revised view of spindle assembly checkpoint activation, MPS1 is recruited to kinetochores marked by high Aurora B phosphorylation. If microtubule attachment is present, MPS1 then triggers destabilization of the attachment and hence formation of an unattached kinetochore, while at the same time promoting recruitment of the components of the spindle assembly checkpoint machinery critical for arresting the cell cycle (Fig. 4H). MPS1 also indirectly promotes the recruitment of the phosphatase PP2A-B56 which is needed to ultimately stabilize microtubule attachments ^25, 26^. This arrangement constitutes an incoherent feed-forward loop with the following stages: MPS1 binds to microtubule-attached kinetochores in an Aurora B-dependent manner and together with Aurora B promotes microtubule release; MPS1 catalyzes spindle checkpoint protein and BUBR1-associated PP2A-B56 recruitment; PP2A-B56 antagonizes the effect of MPS1 and Aurora B on microtubule turn-over and then on MPS1 binding itself; microtubules re-bind and MPS1 leaves kinetochores if the microtubule attachment has sufficiently attenuated Aurora B activity (Fig. 4H). In such a model, the key antagonistic factors for microtubule-kinetochore attachment formation are MPS1 and PP2A-B56. Indeed, a simple epistasis experiment, in which the effect of PP2A-B56 depletion was combined with either Aurora B or MPS1 inhibition, revealed that MPS1 inhibition “rescued” the effect of PP2A-B56 depletion more effectively than Aurora B inhibition, in keeping with the suggested incoherent feed-forward loop (Figure 4E-G).

**Fig. 4.**
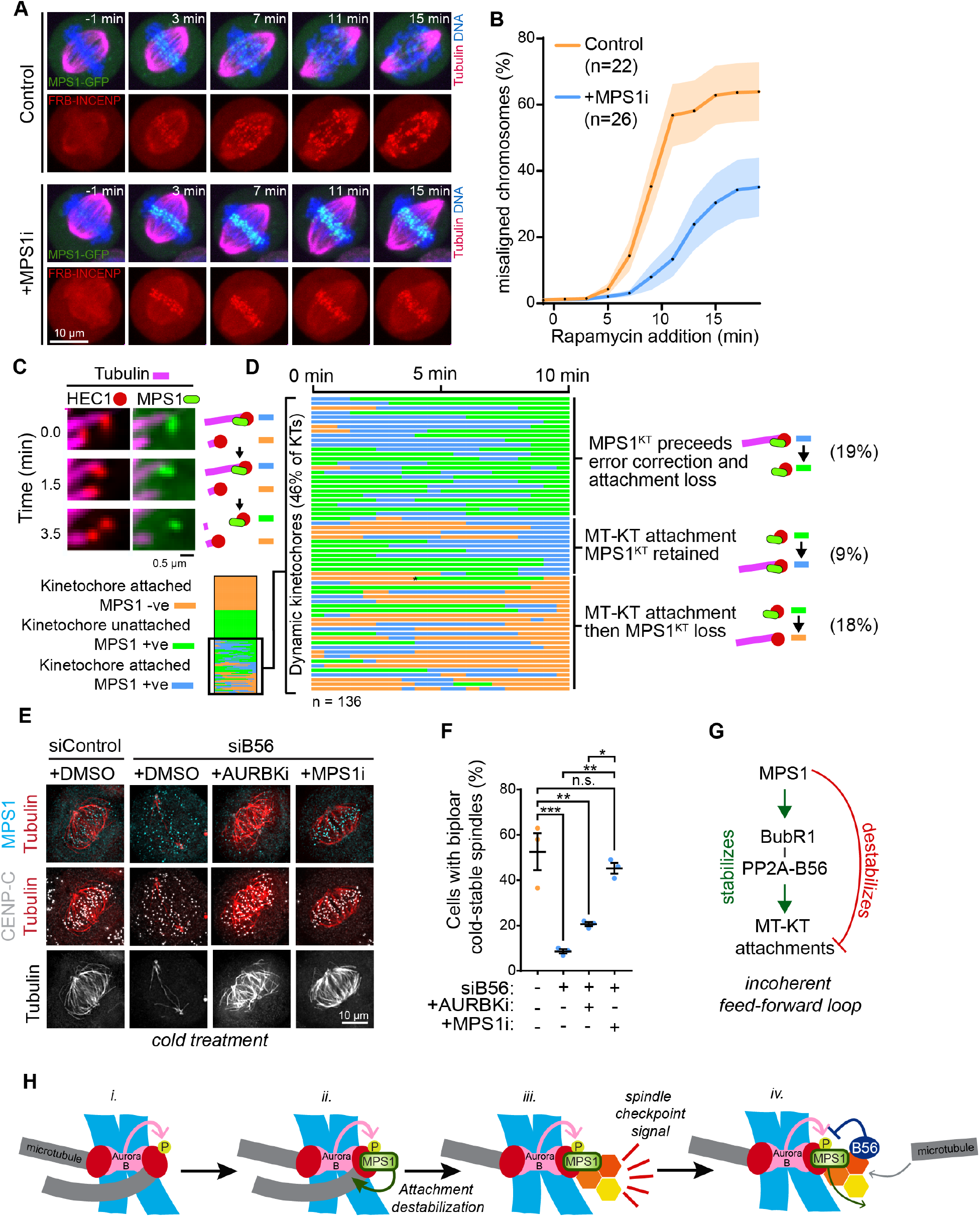
MPS1 generates unattached kinetochores. (**A**) Movie stills of a HeLa cell expressing Myc-Mis12-FKBPx3, mCherry-FRB-INCENP^47-918^ and MPS1-GFP, incubated with SiR-Tubulin (magenta) and Hoechst (blue). Cells were treated with 20 μM MG132 30 min prior to imaging. Cell at -1 min is at metaphase prior to addition of 500 nM rapamycin. MPS1i AZ3146 was added at 2 μM for 10 min prior to imaging. (**B**) Quantification of misaligned chromosomes (blue) in cells as shown in (A). (**C**) Live cell imaging stills of HeLa cells expressing HEC1-mCherry and MPS1-GFP, incubated with SiR-Tubulin and 10 μM STLC prior to imaging. Shown is an example of an initially syntelically attached kinetochore pair, where MPS1-GFP localization precedes error correction and attachment loss at one kinetochore. (**D**) 136 kinetochores of the cells in (C) were assessed for kinetochore attachment status and presence of absence of MPS1-GFP over a 10-minute period. (**E**) Immunofluorescence analysis of control or PP2A-B56 depleted HeLa MPS1-GFP cells treated with the indicated kinase inhibitors before cold-treatment. (**F**) Quantitation of cells treated as in (F), n =3. (**G**) Incoherent feed-forward loop describing MPS1 and PP2A-B56 kinetochore recruitment and effector function. (**H**) Model for MPS1 recruitment and function.

Current thinking is propelled by the notion that a tension-dependent NDC80 phosphorylation-dephosphorylation cycle underpins error correction and checkpoint signaling ^1^. In this model, Aurora B phosphorylates NDC80 at incorrect low-tension attachments, which become destabilized and attract MPS1 to initiate the spindle checkpoint signal. Conversely, microtubule-binding displaces MPS1 and stops the checkpoint signal. Here we report that MPS1 kinetochore binding is not competitive with microtubule binding and is instead recruited to microtubule-attached kinetochores marked by Aurora B activity. This pool of MPS1 then actively triggers microtubule release, generating checkpoint-positive kinetochores. Our data are consistent with findings showing that spindle checkpoint silencing can occur with only 20% kinetochore microtubule occupancy ^15^, consistent with a ‘state’ change of the kinetochore upon microtubule attachment determined by the kinetochore kinase-phosphatase balance, as postulated here. Intriguingly, we and others have also shown that MPS1 recruitment does not depend on Aurora B phosphorylation of the N-terminus of NDC80 ^4^ (Extended Data Fig. 8). Instead, NDC80 phosphorylation by Aurora A appears crucial in a pathway for mitotic spindle size control (Sobajima et al. 2022, bioRxiv). Taken together, these findings suggest there is a need to reevaluate the role of Aurora B in error correction and spindle checkpoint initiation. Further work is required to identify the crucial Aurora B and MPS1 targets, and understand their roles in MPS1 recruitment and regulation of microtubule-binding to the kinetochore, respectively.

Our findings lead to an important revision to our understanding of spindle checkpoint signaling, whereby MPS1 triggers correction of the erroneous microtubule-kinetochore attachment and catalyzes the spindle checkpoint “wait” signal at the same time. Different to current thinking, MPS1 is not a passive sensor for unattached kinetochores, and instead plays a direct role in their generation. Since error correction during microtubule-kinetochore attachment formation and the spindle assembly checkpoint are crucial control mechanisms that safeguard genome stability during chromosome segregation, these insights have important implications for cancer research, where inhibitors of both Aurora B and MPS1 are currently tested in clinical trials.

## Materials and Methods

### Reagents and antibodies

General laboratory chemicals and reagents were obtained from Sigma-Aldrich and ThermoFisher Scientific unless specifically indicated. Inhibitors were obtained from Tocris Bioscience (MPS1 inhibitor AZ3146 (3994); Aurora B Kinase Inhibitor ZM447439 (2458); PP1 and PP2A inhibitor Calyculin A (1336)), Sigma-Aldrich (Kinesin-5/Eg5 inhibitor S-trityl-L-cysteine (STLC) (164739-5G); Rapamycin (R0395)), Santa Cruz Biotechnology (Proteasome inhibitor MG132 (sc-201270)), Spirochrome (SiR-Tubulin (SC002)) and Merck Chemicals Ltd (microtubule polymerization inhibitor Nocodazole (484728-10MG)).

Transfection reagents oligofectamine (Invitrogen) for siRNA oligos, and LT1 (Mirus-Bio) for plasmids were used according to the manufacturers’ instructions. Opti-MEM medium (Gibco) was used to set up transfections.

Commercially available antibodies were used for CENP-C (Guinea Pig pAb; MBL (PD030); 1:2000 dilution for immunofluorescence), BubR1 (Rabbit pAb; Bethyl (A33-386A); 1:1000 dilution for western blotting), Tubulin (Mouse mAb (DM1A); Sigma-Aldrich (T6199); 1:2000 dilution for immunofluorescence, 1:5000 dilution for western blotting), MPS1 (Mouse mAb (N1); Abcam (ab11108); 1:2000 dilution for western blotting), beta Actin (Mouse mAb (AC-15) HRP conjugate; Abcam (ab49900); 1:5000 dilution for western blotting), HEC1 (Mouse mAb (9G3.23); GeneTex (GTX70268); 1:1000 dilution for western blotting). Rabbit antibodies against Astrin and Sheep antibodies against HURP have been described previously ^36, 37^.

All secondary antibodies were used at 1:1000 dilutions based on recommended stock concentrations. Secondary donkey antibodies against mouse, rabbit or sheep and labelled with Alexa Fluor 350, Alexa Fluor 555, Alexa Fluor 647 were purchased from ThermoFisher Scientific. Secondary donkey antibodies against Guinea Pig labelled with Alexa Fluor 647 and secondary donkey antibodies against Mouse, Rabbit or Sheep labelled with HRP were purchased from Jackson ImmunoResearch Laboratories. Protein A and Protein G conjugated to HRP were purchased from Merck. DNA dye Hoechst 33342 was purchased from ThermoFisher Scientific and used at a final concentration of 5 μg/ml. For Western blotting, proteins were separated by SDS-PAGE and transferred to nitrocellulose using a Trans-blot Turbo system (Bio-Rad). Protein concentrations were measured by Bradford assay using Protein Assay Dye Reagent Concentrate (Bio-Rad). All Western blots were revealed using ECL (GE Healthcare).

### Molecular biology

Human MPS1, HEC1 and BUBR1 were amplified from Human testis cDNA (Marathon cDNA; Takara Bio Inc.) using Pfu polymerase (Promega). Mammalian expression constructs were made using pcDNA5/FRT/TO vectors (Invitrogen), modified to encode the EGFP- or mCherry-reading frames. BubR1 mutagenesis has been described previously ^27^ as has generation of MPS1 kinase-dead mutants ^21^.To generate phospho-null (N8A) or phospho-mimetic (N8E) MPS1 autophosphorylation sites, the first 300 base pairs of MPS1 were synthesized by Twist Bioscience with the site encoding amino acids Ser7, Thr12, Ser15, Thr33, Ser37, Thr45, Thr46 and Ser49 changed to encode either alanines (N8A) or glutamic acids (N8E) residues. HiFi assembly (NEB) was then used to replace the first 300 base pairs of MPS1 with the synthetic fragments in expression vectors. The HEC1-9A and 9D ^38^ mutants were generated by gene synthesis and cloned into pCDNA5/FRT/TO encoding a C-terminal GFP-fusion.

### Cell line generation

HeLa cells homozygously expressing MPS1 endogenously tagged with GFP at the C-terminus have been described before ^24^. HeLa cell lines with single integrated copies of the desired transgene were created using the T-Rex doxycycline-inducible Flp-In system (Invitrogen ^39^) using pcDNA5/FRT/TO vectors. MPS1 was endogenously tagged at the N-terminus with GFP using CRISPR/Cas9 editing in HeLa cells already containing a single T-Rex doxycycline-inducible Flp-In integration site using a previously described method ^24, 40^. In brief, homology recombination cassettes containing the desired knock-in DNA with flanking regions of homology of 1075 bp to the target locus were co-transfected with a version of pSpCAS9(BB) (Addgene, 48139) containing the guide RNA sequence 5’-TCTTTGATGCTAGTTAAAGT-3’ and modified to removed puromycin resistance. The knock-in sequences harbor a puromycin resistance marker followed by a glycine-serine rich flexible linker (GS), a P2A ribosome-skipping sequence, and the EGFP protein sequence followed by a glycine-serine rich flexible linker (GS). Antibiotic-resistant clones were selected and successful modification was confirmed by western blotting.

INCENP kinetochore recruitment by rapamycin addition was achieved by editing a previously described system (plasmid pERB109, Addgene number58280) (31). This was adapted by placing miRFKBP5_Mis12-GFP-FKBP3 under doxycycline-inducible expression and replacing GFP with a Myc tag. INCENP 47-918 was inserted after mCh-FRB. 831 and 804 bp homology arms flanking the AAVS1 safe harbour locus were also added, and the system was stably integrated into the AAVS1 safe harbour locus of HeLa cells homozygously expressing MPS1 endogenously tagged with GFP at the C-terminus by CRISPR/Cas9 knock-in, using the guide RNA sequence 5’-GTTAATGTGGCTCTGGTTCT-3’.

### RNAi and RNAi rescue assays

All siRNA depletions were performed for 48 hours. Control siRNA was performed against GL2 (luciferase) (5’-CGUACGCGGAAUACUUCGAUU-3’) (Dharmacon (#D-001100-01-20). siRNA oligos targeting PP2A-B56 have been described previously ^27^ and were purchased from Dharmacon. All other siRNA oligonucleotides were purchased from ThermoFisher Scientific. RNAi rescues assays with GFP-MPS1/mCherry-BUBR1 transgenes were all performed using the same protocol. Transgene induction was initiated with the addition of 2 μM doxycycline (InvivoGen) 2 hours prior to siRNA addition. Endogenous MPS1 was depleted using oligonucleotides against the 3’ UTR (5′-UUGGACUGUUAUACUCUUGAA-3′, 5′-GUGGAU AGCAAGUAUAUUCUA-3′, and 5′-CUUGAAUCCCUGUGGAAAU-3′) ^24^. Endogenous HEC1 was depleted using oligonucleotides against the 5’ UTR (5’-CCCUGGGUCGUGUCAGGAA-3’) ^4^. Endogenous BUBR1 was depleted using oligonucleotides against the 3’ UTR (5′-GCAATC AAGTCTCACAGAT-3′) ^27^. A second induction was performed 24 h into the siRNA depletion.

### Mitotic arrests, kinase inhibitions, rapamycin addition and cold treatments

Nocodazole (0.6 μM) treatments of cells to depolymerize microtubules and arrest the cells in a prometaphase state were performed for 2 hours. STLC (10 μM) treatment to inhibit Eg5 kinesin and generate monopolar spindles was also performed for 2 hours. MG132 (20 μM) to inhibit the proteasome and arrest cells in metaphase was performed for 30 min. MPS1 inhibition was performed using AZ3146 (2 μM) for 15 min. Aurora B inhibition was performed using ZM447439 (10 μM) for 10 min. When cells arrested in nocodazole or STLC were treated with AZ3146 or ZM447439, or in experiments with the MPS1 phospho-mutants, the proteasome inhibitor MG132 (20 μM) was added 30 min prior to fixation to prevent premature mitotic exit. Rapamycin was added at 500 nM, with times indicated in figures. Mitotic cells were cold treated by placing cell dishes on ice and replacing the existing media with media (DMEM with 1% (vol/vol) GlutaMAX (Life Technologies) containing 10% (vol/vol) bovine calf serum) at 4°C. Cells remained on ice for 9 min then were fixed at room temperature.

### Immunofluorescence staining

Cells were fixed with PTEMF buffer (20mM PIPES-KOH pH 6.8, 0.2% v/v Triton X-100, 10mM EDTA, 1mM MgCl_2_, 4% v/v formaldehyde) for 12 minutes at room temperature. Coverslips were washed in phosphate buffered saline (PBS) and incubated in blocking buffer (3% w/v bovine serum albumin, 0.1% v/v Triton X-100 in PBS) for a minimum of 45 minutes. Coverslips were incubated face-down on 80μl droplets of primary antibodies in a humidified chamber for 1 hour. Following primary antibody incubation coverslips were washed 3 x in PBS. Secondary donkey antibodies against mouse, rabbit, guinea pig, or sheep, labelled with Alexa Fluor 405, Alexa Fluor 555, or Alexa Fluor 647 (Molecular Probes) were used at 1:1000. Coverslips were incubated face-down on 80μl droplets of diluted antibodies in a humidified chamber for 45 minutes. Coverslips were washed 3 x in PBS and 1 x in distilled deionised water. Coverslips were left to dry completely before being mounted with 7μl of Mowiol 4-88 (Sigma) according to manufacturer’s instructions. For the images in Fig. 2G, coverslips were mounted onto droplets of Vectashield Plus (2bscientific) and sealed with clear nail polish.

### Immunofluorescence microscopy

Samples seeded on #1.5 thickness coverslips were imaged on a DeltaVision Core light microscopy system (GE Healthcare) using a 100×/1.4-NA objective fitted to an Olympus IX-71 microscope stand. Standard filter sets for DAPI (excitation 390/18, emission 435/48), FITC (excitation 475/28, emission 525/48), TRITC (excitation 542/27, emission 597/45), and Cy-5 (excitation 632/22, emission 676/34) were used to sequentially excite and collect fluorescence images on a CoolSnap HQ2 CCD camera (Photometrics) using the software package softWoRx (GE Healthcare). Cells were imaged using a 0.2-μm interval and a total stack of 2 μm and deconvolved for presentation using softWoRx. For quantification, imaging was performed using a 60×/1.35-NA oil-immersion objective on a BX61 Olympus microscope equipped with filter sets for DAPI, EGFP/Alexa Fluor 488, 555, and 647 (Chroma Technology Corp.), a CoolSNAP HQ2 camera (Roper Scientific), and MetaMorph 7.5 imaging software (GE Healthcare).

### Super-resolution immunofluorescence microscopy

Samples seeded on #1.5 thickness coverslips were imaged on an Olympus SoRa spinning disk confocal microscope using a 60x/1.5-NA objective fitted to an Olympus IX-83 microscope stand with 3.2 x optical zoom and a Yokogawa CSU-W1 SoRa super-resolution spinning disk. Solid state lasers emitting 405 nm, 488 nm, 561 nm and 633 nm were used. Images were captured with a Prime BSI sCMOS camera (photometrics) using Olympus cellSens software package. Images were acquired with a 0.24 μm interval over a total distance of 4.8 μm. Constrained iterative deconvolution was performed on cellSens.

### Live cell imaging

Cells seeded on circular glass bottom Fluorodish imaging dishes (World Precision Instruments) in Fluorobrite media (ThermoFisher Scientific) supplemented with 10% FBS and 1x GlutaMAX (ThermoFisher Scientific) were imaged at 37°C with 5% CO_2_ on an Olympus SoRa spinning disk confocal microscope using a 60x/1.5-NA or 100x/1.45-NA objective fitted to an Olympus IX-83 microscope stand with a Yokogawa CSU-W1 SoRa super-resolution spinning disk. Solid state lasers emitting 405 nm, 488 nm, 561 nm and 633 nm were used. Images were captured with a Prime 95B sCMOS camera (photometrics) using Olympus cellSens software package. Kinetochores of STLC arrested cells were imaged using the 100x/1.45-NA objective as 1.3 μm stacks with intervals of 0.26 μm across a period of 15 minutes with 30 second intervals. STLC (10 μM) was added 4 hours prior to imaging. 10 μm stacks with intervals of 0.5 μm of MG132/Nocodazole arrested cells were imaged using the 60x/1.5-NA objective at intervals of 2 min over a total period of 22 minutes, with 100 ul PBS + Rapamycin added in the interval between the first a second timepoint. SiR-Tubulin (50 nM) and Hoechst (4 μM) were added 1 hour prior to imaging.

### Quantification and statistical analysis

#### Image processing and analysis

Image processing and analysis was performed using the Fiji distribution of ImageJ ^41^. For figures, deconvolved images acquired on a DeltaVision Core light microscopy system were cropped to 350 × 350px and maximum projected. Channels were subject to linear contrast adjustment. Within each figure contrast adjustment of each channel is the same between conditions and merges.

Quantitation was performed on images of cells from a BX61 Olympus microscope which were cropped to 250 × 250px and sum projected through 7 z-slices. Kinetochore intensities for each fluorescence channel were determined by placing 8px-diameter circular ROIs at the maxima of individual non-overlapping kinetochores and measuring the mean pixel intensity of each channel within said selections. Where possible 20 kinetochores were measured per cell. Background measurements were derived by taking an equivalent number of pixels as were in the ROI which were as close as possible to the ROI without overlapping with kinetochores. In brief, a binary mask of kinetochore signal was generated by performing a tophat transform of the CENP-C channel and thresholding using an iterative intermeans method ^42^. Pixels were radially selected from outside the kinetochore ROI and, if not overlapping with signal in the binary kinetochore mask, added to a new background ROI. Once 52px has been incorporated into the background ROI the mean pixel intensity of each channel within said ROI was measured.

Data analysis was performed in Rstudio ^43^ using the Tidyverse collection of packages ^44^. Kinetochore signal intensities were background-adjusted by subtracting the background signal on a channel-by-channel basis. Next, the mean intensity of the channel of interest was divided the mean intensity of the CENP-C channel on a per-kinetochore basis. The mean kinetochore localization intensities were then calculated for each cell. Normalization was performed within repeats by dividing each cell’s mean kinetochore localization intensity by that of the group which was being normalized to.

Measurements of microtubule width, and of HURP and Tubulin intensity on attached K-fibers, were performed in Fiji on DeltaVision acquired images prior to deconvolution. Microtubule width was measured on single Z-planes of metaphase microtubule bundles which were clearly attached to a kinetochore and had no adjacent microtubule bundles. A 1 μm line was drawn from the center of the kinetochore through the axis of the microtubule, then the line was rotated 90° so that the line was perpendicular to the microtubule and 0.5 μm away from the kinetochore. Measurements of pixel intensity along the line were taken. A subsequent 1 μm line was then drawn parallel to the microtubule bundle and the average pixel intensity along that line was subtracted as background signal. Measurements of HURP and Tubulin intensity were taken from projected images stack of STLC or MG132 arrested cells respectively. Measurements were only taken from microtubule bundles which were clearly attached to a kinetochore and had no adjacent microtubule bundles. Measurements were recorded from a circular ROI with a 0.25 μm radius placed 0.5 μm (Tubulin) or 2 μm (HURP) from the edge of the kinetochore at the center of the microtubule bundles. An adjacent cytoplasmic circular ROI was also measured and that value subtracted as background.

Analysis of kinetochore MPS1-GFP and mCherry-FRB-INCENP-47-918 from live imaging was performed by placing 5 8px-diamter circular ROIs over MPS1 positive kinetochores at the first timepoint they were visible. The same ROIs were then measured at previous timepoints. ROIs were moved where needed at subsequent timepoint to track the same kinetochores. Background measurements were taken from 5 chromatin free cytoplasmic regions. Where no MPS1 positive kinetochores were visible, ROIs were taken from 5 points of the mitotic plate, as defined by Hoechst staining. Chromosome misalignment was measured by generating a rectangular ROI at timepoint 0 which covered the metaphase plate as defined by Hoechst staining, and measuring the intensity of Hoechst staining. A second measurement was taken of the total level of Hoechst staining in the cell (the cell boundary defined by areas with MPS1-GFP). A third measurement was made of the average background intensity outside of the cell, which was multiplied by the area of either the metaphase plate ROI of whole cell ROI, and subtracted from those values. A percentage of chromatin outside of the metaphase plate could then be ascertained and reported as percent of chromosomes that are misaligned. This was performed for each timepoint, with the rectangular ROI measuring the area of the metaphase plate moved where needed to be covering what remained of the metaphase plate and so as to be perpendicular with the spindle axis, as defined by SiR-Tubulin staining. Where chromosome misalignment was severe and SiR-Tubulin staining demonstrated that no microtubules remained in the vicinity of the area previously occupied by the metaphase plate, the level of chromosome misalignment was set as 100%.

### Statistical Analysis

All statistical analysis was performed using GraphPad Prism version 9.2.0 for Windows (GraphPad Software, San Diego, California USA, www.graphpad.com). Each cell measured was considered as a biological replicate (n), hence mean measurements calculated for each cell were used for statistical analysis. At least three independent repeats of each experiment were performed, with statistical analysis performed using a sum of biological replicates from all independent experiments.

Data sets were tested for normal distribution via a D’Agostino-Pearson omnibus K2 test. If all groups in an experiment were normally distributed, then the means were compared using a parametric statistical test as follows. If only two groups were compared then an unpaired 2 tailed t-test was used (with Welch’s correction if the groups had unequal standard deviations). If more than two groups were compared with equal standard deviations a one-way ANOVA was used, which if rejected was followed by a Tuckey’s multiple comparisons test. If more than two groups were compared with unequal standard deviations a Brown-Forsythe ANOVA was used, which if rejected was followed by a Dunn’s multiple comparisons test.

If all groups did not exhibit a normal distribution then medians were compared using a non-parametric statistical test as follows. If only two groups were compared then a Mann-Whitney test was used. If more than two groups were compared then a Kruskal-Wallis test was used, which if rejected was followed by a Dunn’s multiple comparisons test. Graphs display the mean ± SEM. p-values are shown on graphs as follows: p > 0.05 = not significant (n.s.), p ≤ 0.05 = *, p ≤ 0.01 = **, p ≤ 0.001 = ***, p ≤ 0.0001 = ****.

## Acknowledgements

We thank Francis Barr for his encouragement, advice, and many fruitful discussions; and Bela Novak and Tomoaki Sobajima for helpful comments on the manuscript.

## Funding

Cancer Research UK Discovery Programme grant DRCNPG-Nov21\100004 (UG, DH)

Medical Research Council grant MR/K006703/1 (UG, DH)

Edward Penley Abraham grant RF 280 (UG, DH)

Medical Research Council studentship (ER)

## Author contributions

Conceptualization: UG, DH, ER

Methodology: UG, DH, ER

Investigation: DH, ER

Visualization: UG, DH, ER

Funding acquisition: UG

Project administration: UG

Supervision: UG

Writing – original draft: DH, UG

Writing – review & editing: UG, DH, ER

## Competing interests

Authors declare that they have no competing interests.

**Extended Data Fig. 1.**
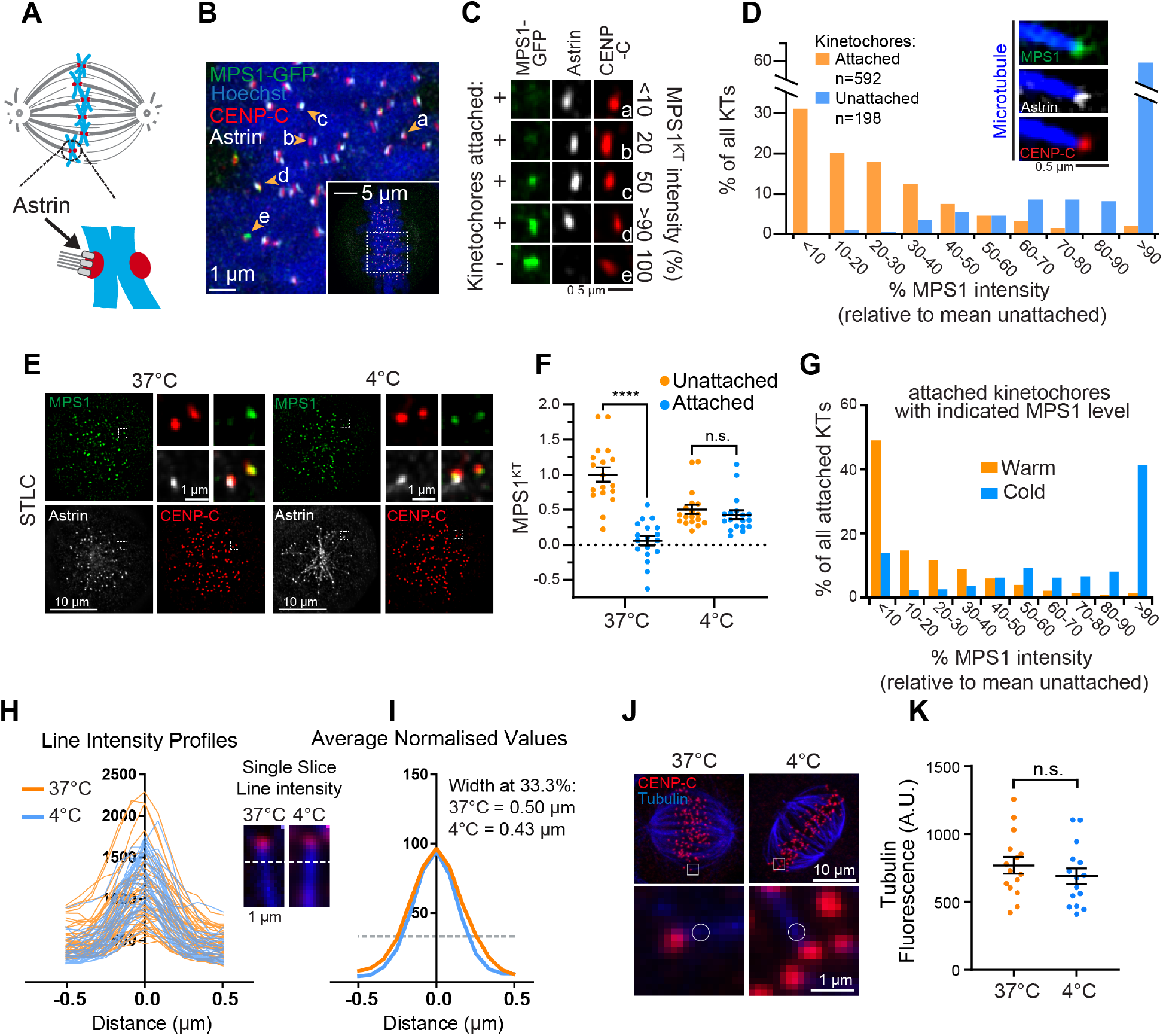
MPS1 can be observed at fully attached kinetochores. (**A**) Diagram of a late prometaphase cell with a monotelically attached chromosome. (**B**) A late prometaphase HeLa MPS1-GFP cell, stained as indicated. Arrows with letters indicate kinetochores displayed in (C). (**C**) Attached and unattached kinetochores from (B) with varying levels of MPS1. (**D**) Histogram of attached kinetochores (Astrin +ve) and unattached kinetochores in prometaphase cells with the indicated MPS1-GFP intensities relative to MPS1-GFP levels at unattached kinetochores. The inset shows an example of an attached kinetochores with high levels of GFP-MPS1. K-fibers were visualized by anti-HURP staining. (**E**) STLC-arrested HeLa MPS1-GFP-cells were either fixed directly or chilled for 9 mins before fixation and staining with the indicated antibodies. (**F**) Quantitation of MPS1 kinetochore intensities from (E), normalized to the mean unattached signal for the given temperature. (**G**) Histogram of MPS1 intensities at attached kinetochores in the warm or cold. (**H**) Raw tubulin intensities (A.U.) across a 1 μm line drawn 0.5 μm from the kinetochore center and perpendicular to the microtubule K-fiber in metaphase cells at 37°C or chilled for 9 min. Example immunofluorescence images are shown with Tubulin (blue) and CENP-C (red). (**I**) Average normalized line profiles from (H) with values at 0.0 μm normalized to 100%. (**J**) Metaphase HeLa cells immunostained for tubulin (blue) and CENP-C (red) at 37°C or chilled to 4°C for 9 min. (**K**) Measurements of tubulin intensity at K-fibers taken 0.5 μm away from kinetochores as shown in (**I**). Error bars represent S.E.M.

**Extended Data Fig. 2.**
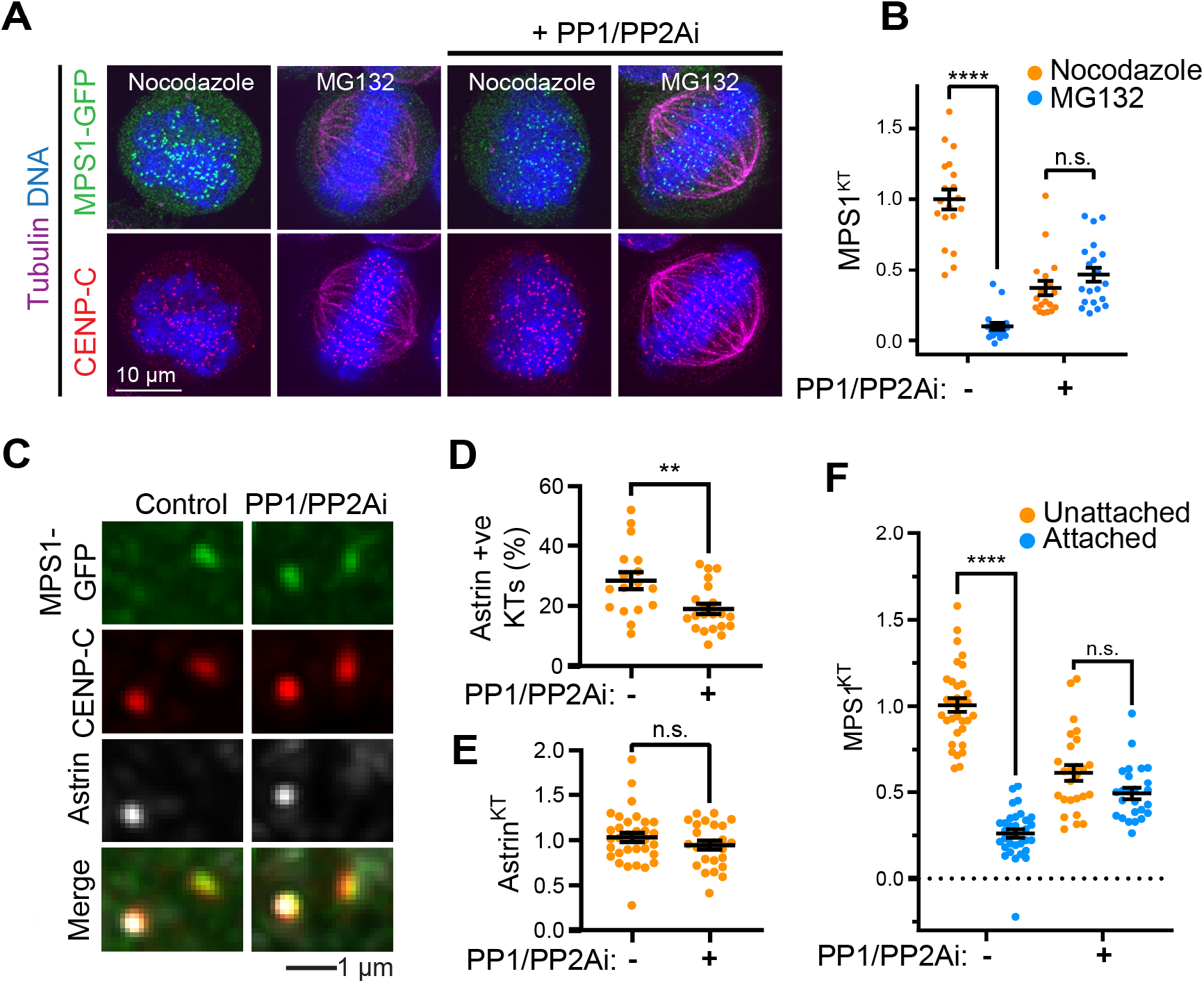
PP1/2A inhibitor treatment retains MPS1 at attached kinetochores. (**A**) Extended version of Figure 2A. HeLa cells with endogenously tagged MPS1-GFP were arrested for 2 hrs with either 0.6μM Nocodazole or 20μM MG132 and then treated with 25 nM calyculin (PP1/2Ai) for 6 min. The cells were immuno-stained as indicated. (**B**) Cell averages of kinetochore-MPS1 at attached metaphase kinetochores in cells treated as in (A). Bars represent mean± S.E.M. Values were normalized to levels of MPS1 at unattached kinetochores without calyculin A treatment. (**C**) STLC arrested HeLa MPS1-GFP cells treated with calyculin A were immuno-stained as indicated. Representative monotelic kinetochore pairs are shown. (**D**) The proportion of attached kinetochores per cell, as determined by the presence of astrin staining, in cells treated as in (C), is plotted. Bars indicate mean± S.E.M. (**E**) Average cell astrin intensities at attached kinetochores in cells treated as in (C) are plotted. Bars show mean± S.E.M. (**F**) Cell average intensities of kinetochore-MPS1 at unattached and attached kinetochores in cells treated as in (C) were plotted. Values were normalized to the mean of the unattached control. Bars show mean± S.E.M.

**Extended Data Fig. 3.**
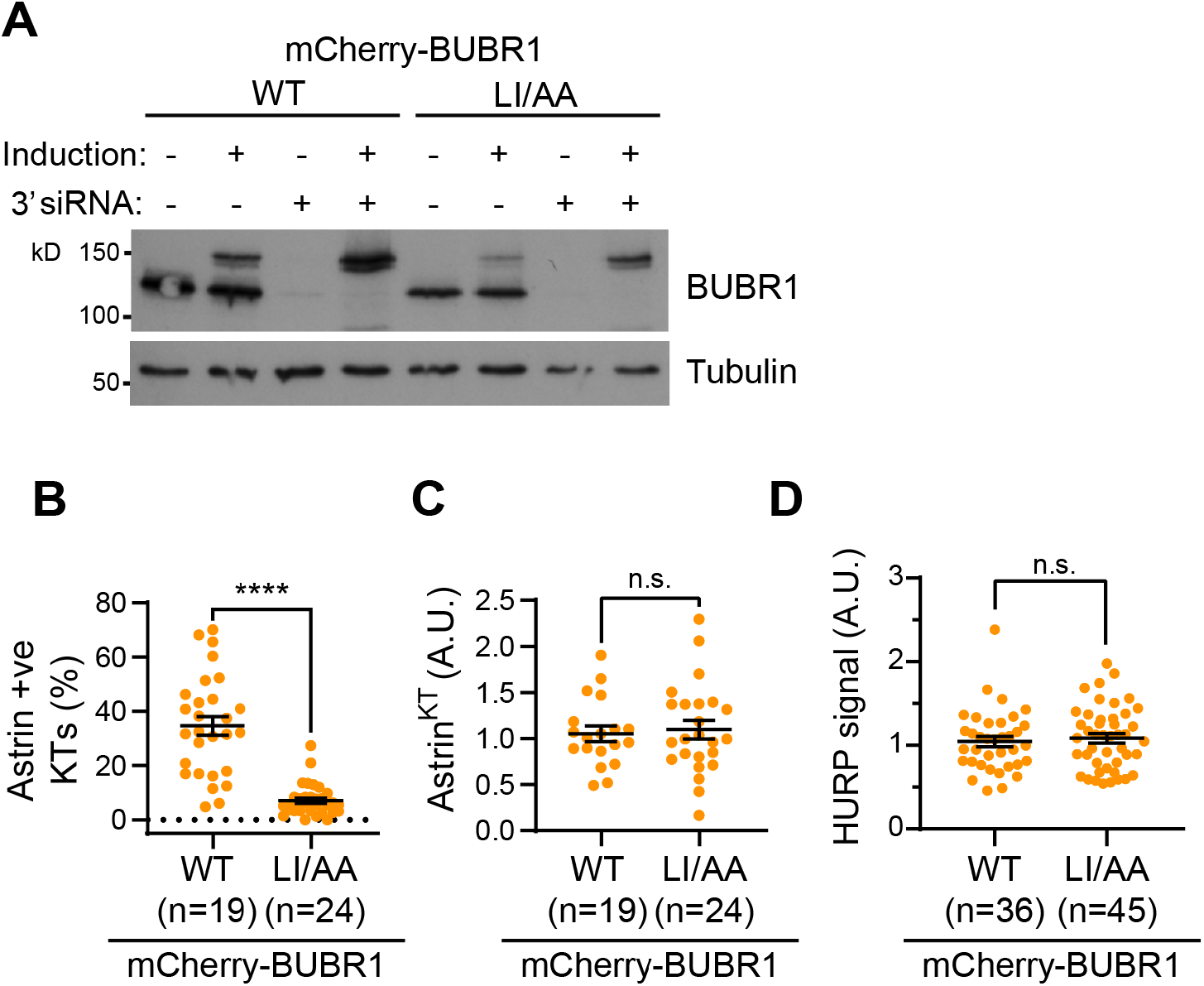
The checkpoint-dependent pool of PP2A-B56 controls the number but not the integrity of microtubule-kinetochore attachments. (**A**) Representative BUBR1 western blot of HeLa TRex-Flp-In cells expressing mCherry-BUBR1^WT^ or mCherry-BUBR1^LI/AA^ demonstrating the depletion of endogenous BUBR1 and induction of the transgenes for the cells used in Fig. 1C. Tubulin is used as a loading control. (**B**) The proportion of attached (Astrin +ve) kinetochores, in cells treated as in Fig. 1C, is plotted. (**C**) Cell average Astrin intensities at attached kinetochores in cells treated as in Fig. 1C were plotted. Bars show mean± S.E.M. (**D**) cell average of measurements of K-fiber HURP intensities in STLC arrested cells treated as in Fig. 1C. Values were normalized to mean of the WT control.

**Extended Data Fig. 4.**
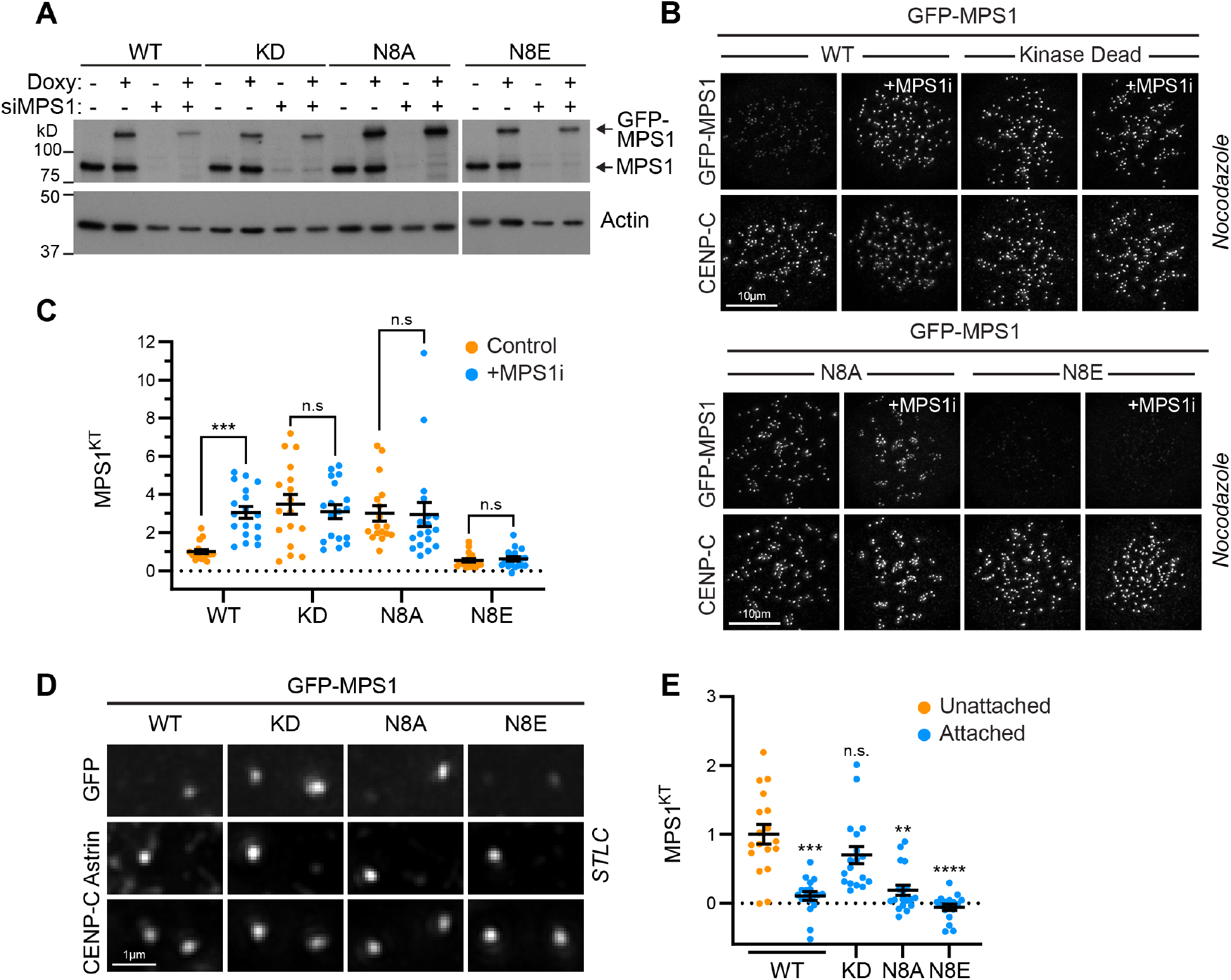
N-terminal autophosphorylation controls the steady-state level of MPS1 at kinetochores. (**A**) Western blot of Flp-In TRex HeLa cells harboring the indicated GFP-MPS1 constructs treated with (+) or without (-) doxycycline induction and with siMPS1 (+) or siControl (-) for 48 hours. Actin was used as a loading control. (**B**) Flp-In Trex HeLa cells depleted of endogenous MPS1 and induced to express the indicated GFP-MPS1 mutant were arrested with nocodazole (2hrs) and MG132 (30min). Cells were treated with or without MPS1 inhibitor AZ3146 (MPS1i) for 15min and immuno-stained as indicated. (**C**) Mean cell averages from (B) of MPS1 kinetochore intensity normalized to mean of untreated GFP-MPS1^WT^ cells are plotted. Bars show mean ± S.E.M. (**D**) Monotelic kinetochore pairs from Flp-In Trex HeLa cells depleted of endogenous MPS1 and induced to express the indicated GFP-MPS1 mutants, and arrested with a monopolar spindle using STLC (2hrs) and MG132 (30mins). (**E**) Cell averages of MPS1 kinetochore intensity at unattached or attached kinetochores from cells in (D), normalized to MPS1^WT^ intensity at unattached kinetochores. Bars show mean ± S.E.M.

**Extended Data Fig. 5.**
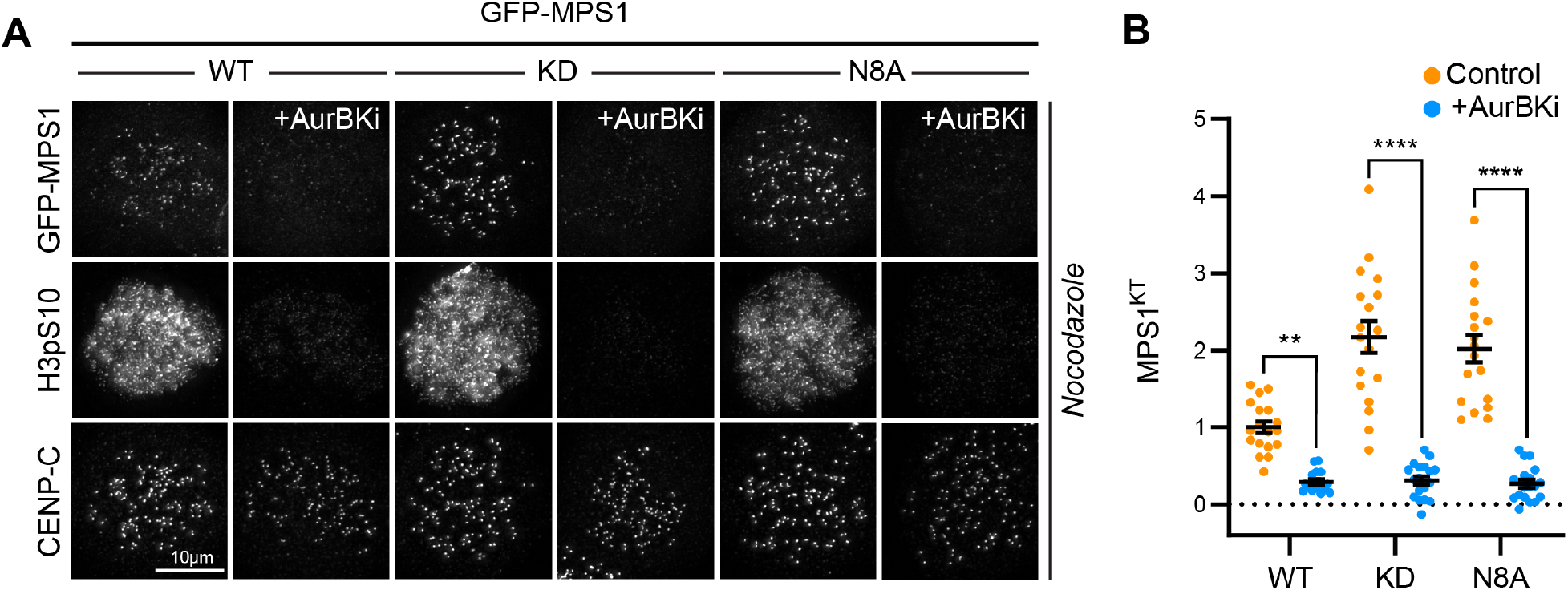
An MPS1 autophosphorylation-site mutant is dependent on Aurora B for localization. (**A**) Flp-In TRex HeLa cells depleted of endogenous MPS1 and induced to express the indicated GFP-MPS1 mutants (KD; kinase dead) were arrested with nocodazole (2hrs) and MG132 (30min). Cells were treated with or without the Aurora B inhibitor ZM447439 (AurBKi) for 10 min and immuno-stained as indicated. (**B**) Cell averages of MPS1 kinetochore intensity from cells in (A). Bars show mean ± S.E.M.

**Extended Data Fig. 6.**
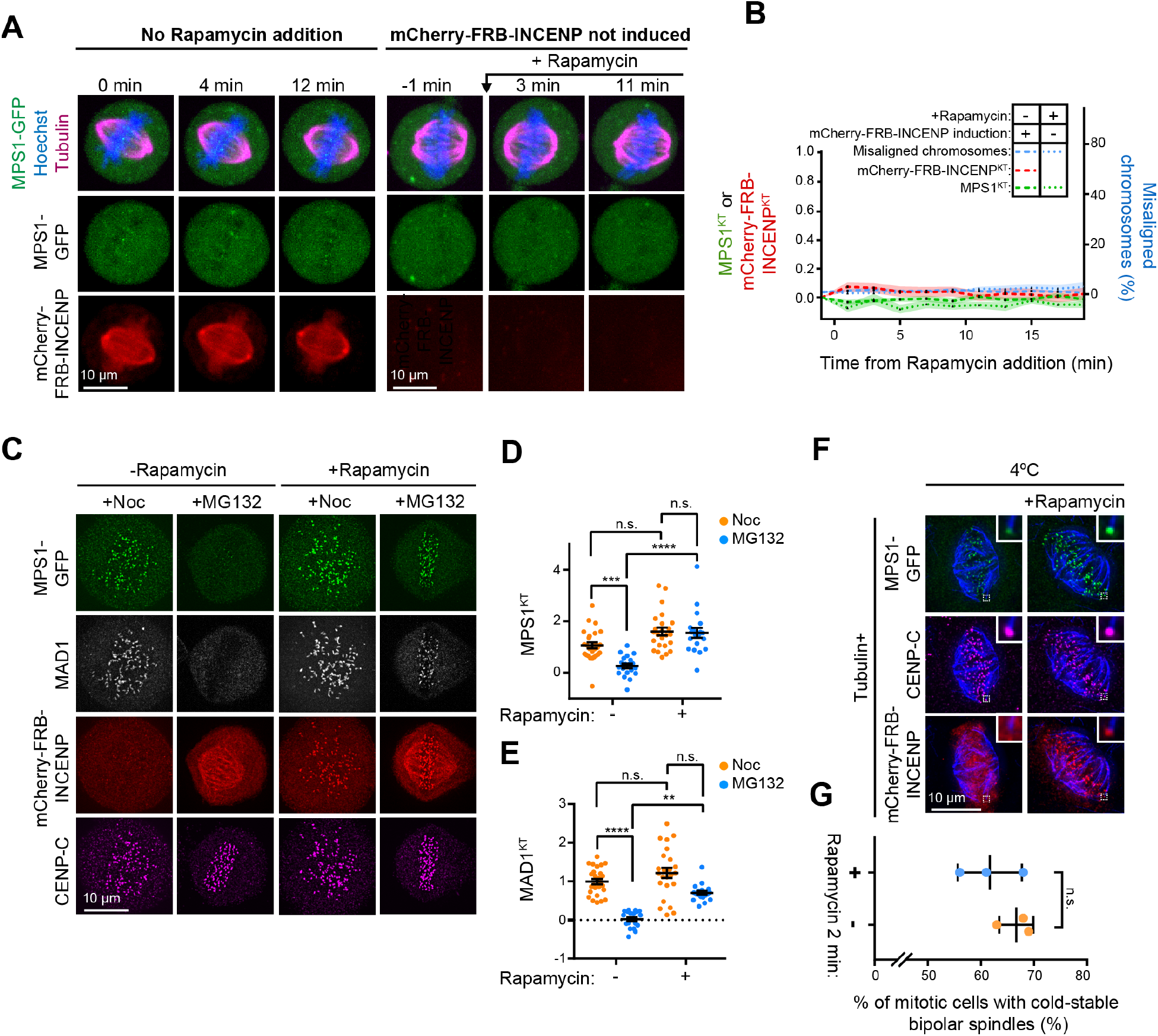
Targeting of INCENP-Aurora B to attached kinetochores results in the recruitment of MPS1 prior to loss of microtubule attachments. (**A**) Live cell imaging stills of HeLa cells expressing MPS1-GFP and incubated with SiR-Tubulin and Hoechst, either with the Aurora-B kinetochore targeting system but without addition of Rapamycin (left), or without the Aurora-B kinetochore targeting system but with Rapamycin added. Cells were treated with 20μM MG132 30 min prior to imaging. Cell at - 1 min is at metaphase before addition of 500nM Rapamycin. In neither case is MPS1-GFP recruited to attached kinetochores, nor is chromosome alignment altered. (**B**) Quantification of kinetochore MPS1-GFP or mCherry-FRB-INCENP intensities and level of chromosome misalignment in cells treated as in (A). Lines represent mean, coloured bands represent S.E.M. (**C**) HeLa MPS1-GFP cells with the Aurora-B kinetochore targeting system were arrested with 0.6 μM Nocodazole or 20μm MG132 for 2 hrs, and 500 nM Rapamycin was added for 2 min prior to fixation. The cells were immuno-stained as indicated. (**D**) Plot of MPS1 kinetochore mean cell intensities from cells in (C). Bars show mean± S.E.M. (**E**) Plot of MAD1 kinetochore mean cell intensities from cells in (C). Bars show mean± S.E.M. (**F**) HeLa MPS1-GFP cells with the Aurora-B kinetochore targeting system were treated as in (C) and then cooled to 4°C for 9 min before fixation and staining as indicated. (**G**) The proportion of mitotic cells with cold-stable bipolar spindles in each condition from 3 experiments is shown. Error bars show S.D.

**Extended Data Fig. 7.**
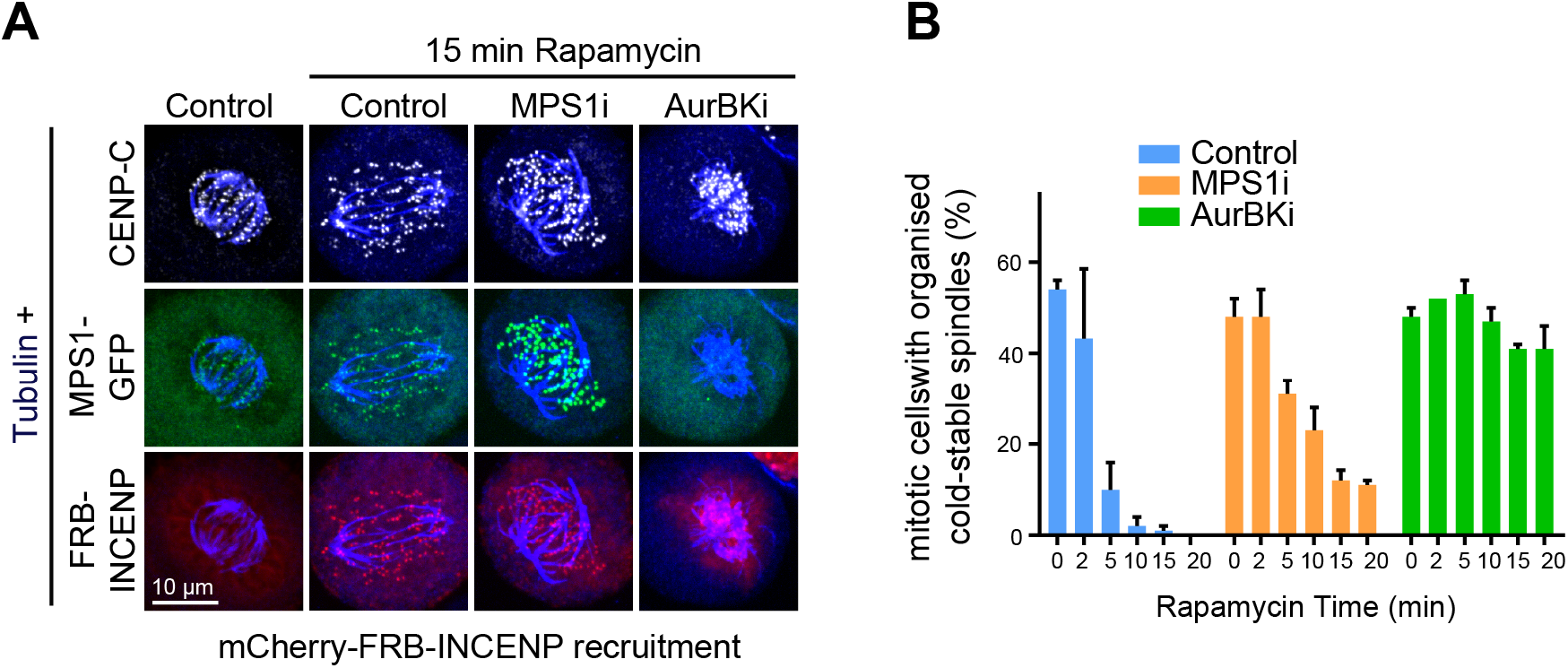
Forced targeting of INCENP-Aurora B and MPS1 to attached kinetochores results in the progressive loss of microtubule attachments. (**A**) HeLa MPS1-GFP cells expressing the Aurora-B kinetochore targeting system were arrested with 20 μm MG132 for 2 hrs, and 500 nM Rapamycin, in the presence or absence of the indicated kinase inhibitors, was added for the indicated time prior to a 9 mins cold treatment followed by fixation and immuno-staining as indicated. (**B**) The proportion of mitotic cells with cold-stable bipolar spindles in each condition at different times after rapamycin addition is shown.

**Extended Data Fig. 8.**
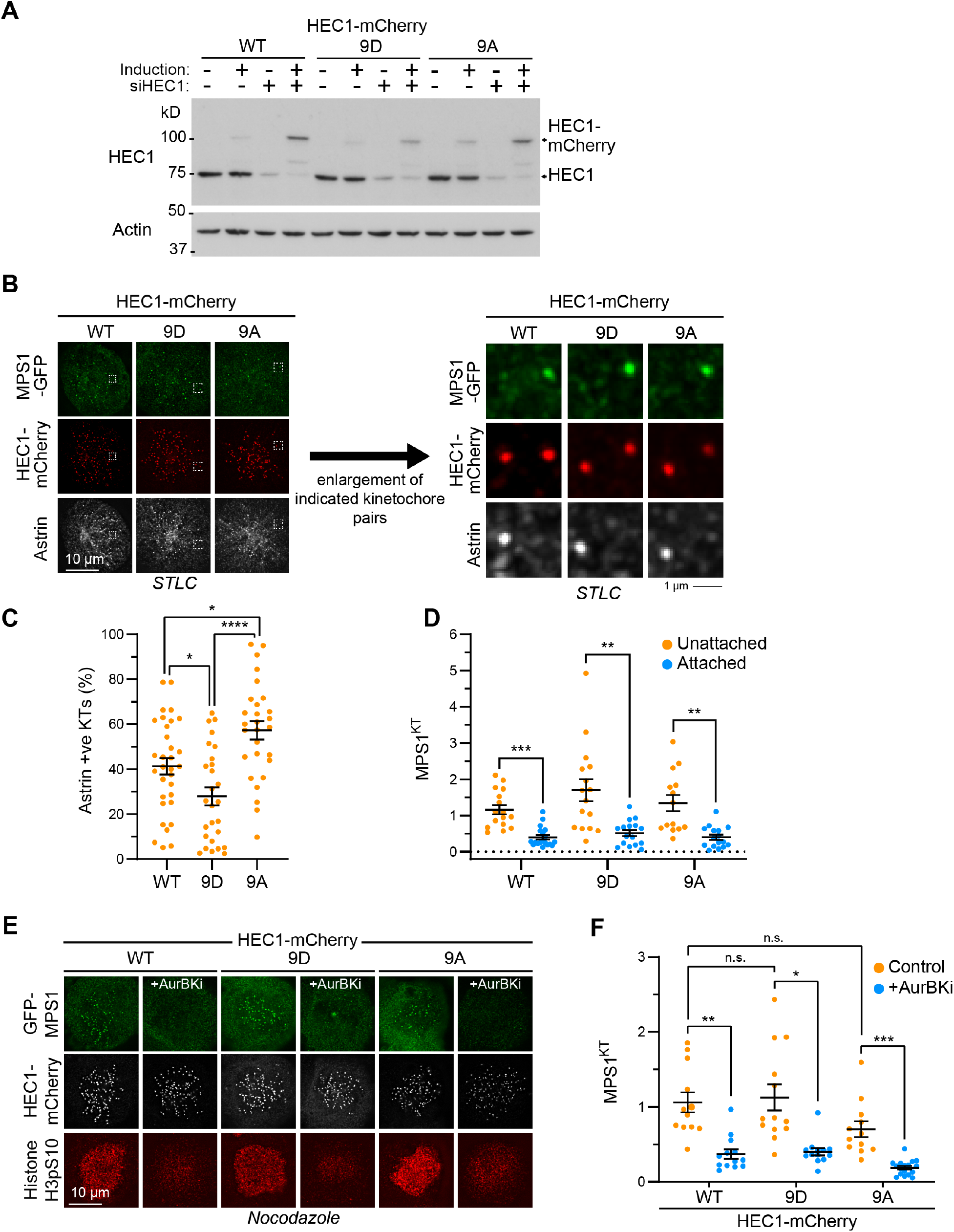
Mutation of HEC1/NDC80-N-terminal phosphorylation sites does not alter MPS1 targeting to kinetochores. (**A**) HeLa Flp-In TRex cells expressing different C-terminally mCherry-tagged phospho-mutants of HEC1, as well as MPS1 endogenously GFP-tagged at the N-terminus, were depleted of the endogenous HEC1 protein as indicated and induced to express the transgene. Cell lysates were prepared and immunoblotted as indicated. Actin was used as a loading control. (**B)** STLC arrested HeLa Flp-In TRex cells, also expressing endogenous GFP-MPS1, were depleted of endogenous HEC1 and induced to express HEC1-mCherry^WT^, HEC1-mCherry^9A^, or HEC1-mCherry^9D^ and immuno-stained with antibodies against Astrin. Representative monotelic kinetochores pairs, indicated by dashed boxes, have been enlarged on the right-hand side. The different HEC1 phospho-mutants do not alter the pattern of MPS1 binding to kinetochores. (**C**) Quantitation of attached (Astrin +ve) kinetochores, in cells treated as in (B), from three independent experiments. Bars represent mean ±S.E.M. (**D**) Cell averages of kinetochore-MPS1 intensities from cells in (B) relative to kinetochore-HEC1 at unattached and attached kinetochores. Values were normalised to mean of the unattached WT condition. Bars show mean ±S.E.M. (**E**) Nocodazole arrested Flp-In TRex HeLa cells, also expressing endogenous GFP-MPS1, were depleted of endogenous HEC1 and induced to express HEC1-mCherry^WT^, HEC1-mCherry^9A^, or HEC1-mCherry^9D^ and treated with DMSO or Aurora B inhibitor ZM447439 (AurBKi) at 10μM for 10 mins. Cells were fixed and stained as indicated. Note that expression of HEC1-mCherry^9A^ still allows MPS1 kinetochore lcalization, and that in cells expressing HEC1-mCherry^9D^ MPS1 is still Aurora B-sensitive. (**F**) Cell average MPS1 kinetochore intensities for the cells in (E) were plotted. Bars indicate mean ±S.E.M.

